# Development of RNAseq methodologies to profile the *in vivo* transcriptome of *Bordetella pertussis* during murine lung infection

**DOI:** 10.1101/140632

**Authors:** Ting Wong, Jesse Hall, Dylan Boehm, Mariette Barbier, F. Heath Damron

## Abstract

*Bordetella pertussis* is an obligate human respiratory pathogen that causes the disease whooping cough. A whole cell vaccine (DTP) was developed in the 1940s and was subsequently replaced in the 1990s with a protein-based subunit acellular vaccine (DTaP; tdap). Today, we are observing a resurgence of whooping cough due to evolution of the pathogen and waning vaccine immunity. The use of vaccines decreased the need for basic research on this pathogen. As a result, numerous questions on the basic pathogenesis of *B. pertussis* remain to be answered. Microarrays and more recently, RNA sequencing (RNAseq), have allowed the field to describe the *in vitro* gene expression profiles of the pathogen growing in both virulent and avirulent phases; however, no published studies have described an *in vivo* transcriptome of the pathogen. To address this need, we have designed and evaluated workflows to characterize the *in vivo* transcriptome of *B. pertussis* during infection of the murine lung. During our initial studies, we observed that only 0.014% of the ~100 million 2x50bp illumina reads corresponded to the pathogen, which is insufficient for analysis. Therefore, we developed a simple protocol to filter the bacteria out of the tissue homogenates and separate bacterial cells from the host tissue. RNA is then prepared, quantified, and the *B. pertussis* to host RNA ratio is determined. Here, we present the protocol and discuss the uses and next directions for which this RNAseq workflow can be applied. With this strategy we plan to fully characterize the *B. pertussis* transcriptome when the pathogen is infecting the murine lung in order to identify expressed genes that encode potential new vaccine antigens that will facilitate the development of the next generation of pertussis vaccines.

## Introduction

*Bordetella pertussis* is a bacterial respiratory pathogen and the causative agent of whopping cough or pertussis. While whopping cough is a vaccine-preventable disease, the number of cases have risen during the past decade (1). Re-emergence of pertussis is a multi-factorial problem that highlights the need to re-design and improve current vaccination strategies. Historically, whole cell vaccines (WCV), made of formalin inactivated bacteria were first introduced in the 1940s. Following implementation of WCV, efforts by multiple research teams led to the identification of *B. pertussis* major virulence factors, the adenylate cyclase toxin (ACT) and pertussis toxin (PT) (2). The global regulatory system of virulence genes *(Bordetella* virulence genes system or Bvg) was discovered (3, 4) and important surface adhesins such as the filamentous hemagglutinin (FHA), fimbriae, and pertactin (PRN) were identified. This knowledge led to the development of acellular vaccines which are composed of PT, FHA, fimbriae, and PRN (5). Since the implementation of acellular vaccination against pertussis, research efforts on the pathogenesis and basic bacteriology of *Bordetella pertussis* have decreased. As a result, there are still very important gaps in the knowledge of how the bacterium causes infection. These gaps need to be filled to rationally design the next generation of pertussis vaccines.

Recent advances in sequencing technology have opened new doors to study bacterial pathogenesis using a global transcriptomics approach. In a previous study, we demonstrated the feasibility of sequencing bacterial RNA in complex samples (6). We were able to isolate both bacterial and host RNA during an acute murine pneumonia, and perform dual-seq to determine both transcriptional profiles during infection. In the present study, we hypothesized that this methodology could be applied for the study of other bacterial pathogens such as *Bordetella pertussis*. Here, we describe the first transcriptome of *B. pertussis* during lung infection and identify some of the technical pitfalls of this approach for tissues in which the bacterial burden is low. To circumvent these technical difficulties, we propose a novel methodology to facilitate bacterial RNA recovery from these samples. We expect that this method will be broadly applicable for the study of the transcriptome of bacterial pathogens during infection.

## Materials and Methods

### Bacterial strains and growth conditions

*B. pertussis* strain UT25 (UT25Sm1) (7) was cultured on Bordet Gengou (BG) agar (8) (remel) supplemented with 15% defibrinated sheep blood (Hemostat Laboratories) for 48 h at 36°C. *B. pertussis* was then transferred from BG plates to three flasks of 12 ml of modified Stainer-Scholte liquid medium (SSM) (9). SSM cultures were not supplemented with cyclodextrin (Heptakis(2,6-di-O-methyl)-β-cyclodextrin). SSM cultures were grown for ~22 h at 36°C with shaking at 180 rpm until the OD_600_ reached 0.5 on a 1 cm path width spectrophotometer (Beckman Coulter DU 530). The cultures were then diluted to provide a challenge dose of 2 × 10^7^ CFU in 20 μl. For growth of *Pseudomonas aeruginosa* strain PAO1, *Pseudomonas* Isolation Agar (Difco) was used. *Bordetella bronchiseptica* RB50 and *Escherichia coli* TOP10 were cultured on Lysogeny Agar (10 g NaCl, 5 g Yeast Extract, 10 g tryptone). *P. aeruginosa, B. bronchiseptica*, and *E. coli* were cultured at 36°C for 18 hr.

### Murine *B. pertussis* challenge

Outbred CD1 mice were obtained from Charles River and NSG mice (NOD.Cg-Prkdcscid Il2rgtm1Wjl/SzJ; Jackson Labs stock number 005557) raised in house by the WVU Transgenic Animal Core Facility. Mice were anesthetized by intraperitoneal injection of ketamine and xyalzine in saline. Two 10 μl doses of the *B. pertussis* strain were pipetted directly into each nostril of the mouse. Five to seven mice were infected with strains UT25, and at 1 and 3 days post challenge, mice were euthanized for determination of bacterial burden in the nasal wash, trachea, and lungs. To determine the number of *B. pertussis* in the nares, 1 ml of PBS was flushed up through the nares and collected. Trachea and lungs were extracted and homogenized. Serial dilutions in PBS were plated on BG containing streptomycin (100 μg/ml) to ensure that only UT25 *B. pertussis* were cultured. All murine infection experiments were performed according to protocols approved by the West Virginia University Institutional Animal Care and Use Committee (IACUC) protocol number 14–1211), conforming to AAALAC International accreditation guidelines.

### Isolation of RNA, library construction, and Illumina sequencing

For the RNA sequencing experiment described in Fig. 2 and 3 of this study, RNA was isolated using RNeasy Mini Kit (Qiagen) as specified by the instructions of the manufacturer. The resulting RNA was treated with RNase-free DNase (Qiagen). To remove the DNase, the samples were then cleaned up on another RNeasy Mini column. The resulting RNA was quantified on a Qubit 3.0 fluorometer (ThermoFisher). Next, the RNA integrity was assessed by running the samples on an Agilent BioAnalyzer RNA Pico chip. Due to the fact that lysis of lung tissue causes RNA degradation and that the RNA has both bacterial and murine ribosomal RNA, we observed low RIN numbers of 2–4. However, there was sufficient RNA from 200 to 1000 nucleotides to prepare libraries. Overall, all of the samples were observed to have similar RIN numbers. For comparison with *in vitro* grown *B. pertussis*, we used samples from SSM *in vitro* grown UT25 extracted and analyzed by RNAseq using an identical procedure previously published by our laboratory (10). These *In vitro* grown *B. pertussis* UT25 RNA samples had RIN numbers of 9–10. For library preparation, illumina Scriptseq complete gold (epidemiology; for mouse/bacteria ribosomal depletion) was used. Resulting libraries passed standard illumina quality control PCR and were sequenced on an illumina Hiseq 1500 by the Marshall University Genomics Core. Three biological samples were pooled into 3 technical samples (9 total mice).

### RNAseq and bioinformatics analyses

The reads were aligned to the *B. pertussis* Tohama I genome (11) using CLC Genomics Workbench version 9.5. Reads per kilobase per million (RPKM) and fold change for each gene were calculated. Approximately 180,000–200,000 reads mapped per 100 million 2x50bp samples. Upon inspection of the reads, we observed cross mapping reads that are likely due to the presence of residual murine RNA in the sample. To avoid non-specific mapping and remove these reads, we increased mapping stringency to 100% identity and extracted only reads that mapped as pairs and counted them as fragments. These manipulations removed ~90% of the total reads. Due to the low number of total reads obtained from the lungs of infected mice, we sampled *in vitro* grown RNAseq reads from 10 million 2x50bp down to 15,000 total reads. Fold change was calculated by comparison of the *in vitro* UT25 *B. pertussis* reads to the *in vivo* NSG mouse UT25 reads. The raw data reads will soon be submitted to the Sequence Read Archive (SRA). If requested, we would also be happy to provide the raw reads directly.

### Filtration of *B. pertussis* from culture and infected murine lungs

To determine the percent recovery of bacteria from *in vitro* cultures, *B. pertussis, B. bronchiseptica, E. coli*, and *P. aeruginosa* were grown in the culture conditions described above. Various amounts from OD_600_ 0.3 to OD_600_ 0.003 were filtered through 5 μm syringe filters (Sartorius Minisart part ref 17594 Cellulose Acetate). The input culture and outflow were plated on agar plates (BG or LA) to determine the amount of input and flow through bacteria. To isolate bacteria from the infected mouse lungs, mice were dissected and the lungs were extracted and placed into 1 ml of PBS. The lungs were then homogenized with glass Dounce tissue grinders (Sigma-Aldrich). The homogenate was then strained through a 70 μm nylon cell strainer (VWR). The filtrate was then passed through a 5 μm syringe filter. The suspension was next pelleted by 16,100 x g for 4 min in 1.5 ml Eppendorf tubes. The supernatant was then discarded, RNAprotect bacterial reagent (Qiagen) was added to the pellet, and samples were stored at −80°C.

### RNA isolation and reverse transcriptase PCR (qRT-PCR)

For qRT-PCR analysis, cells stored at −80°C in RNAprotect (Qiagen) were lysed with lysozyme, and RNA was isolated using the RNAsnap method (12). Briefly, cell pellet were resuspended in RNA extraction solution (18mM EDTA, 0.025% SDS, 1% 2-mercaptoehanol, 95% formamide) by vortexing vigorously. Samples were incubate at 95°C for 7 min and pelleted by centrifugation at 16,000g for 5 min at room temperature. The supernatant containing RNA was then pipetted into a fresh tube without disturbing the clear gelatinous pellet, and DNA and RNA concentrations were measured using a Qubit 3.0 fluorometer. After extraction, DNA was digested using an off-column digestion with RNase-free DNase and re-isolated with another RNeasy column. RNA concentration and quality was assessed on a Molecular Devices i3 Spectramax Spectra drop plate and on a Qubit 3.0 fluorometer. To ensure RNA was DNA-free, 25 ng of RNA was checked by PCR amplification and was only used for cDNA if no amplicon was observed and a C_T_ of >32. cDNA was synthesized using M-MLV reverse transcriptase (Promega) per the manufacturer’s instructions using 200 ng of RNA and gene specific reverse primers for targets. Twenty five microliter qPCR mixtures were setup with Excella SYBR Green PCR master mix (Worldwide Medical Products), per manufacturer’s instructions using 1 μl of cDNA. A minimum of three technical replicate reactions were ran per gene target per sample on a Step One Plus qPCR thermocycler (Applied Biosystems). Primers were designed on Primer3 (Primer-Blast; NCBI) and checked for specificity by PCR. Melt curve analysis as well as subsequent agarose gel electrophoresis were performed on all reactions. Gene expression was normalized to the *rpoB* reference using the 2^−ΔΔC_T_^ method (13). For statistical analysis, the △Ct for the three biological replicate experiments was calculated and a Student’s *t-test* was performed using Microsoft Excel 2013. Standard error of mean was calculated based on the variability of the △Ct of three biological replicates. For the *rpoB* to *Gapdh* absolute quantitation PCR, amplicons of *rpoB* and *Gapdh* were generated, purified and quantified. The number of copies per ng of DNA was determined and standard curves were generated for *rpoB* (5x10^8^ copies to 500) and *Gapdh* (4.4×10^8^ to 440 copies) with three technical replicates. Microsoft Excel was used to plot standard curves based on the CTs of the known standards described above and number of copies in each sample was calculated using a linear regression with R^2^ values of 0.999 for both *rpoB* and *Gapdh*. The ratio of *Gapdh* to *rpoB* RNA was used to estimate the relative amount of host to pathogen RNA in each sample. The following primers sequences used in this study were described by Bibova *et al*. (14): cyaF(CGAGGCGGTCAAGGTGAT), cyaR(GCGGAAGTTGGACAGATGC), ptxAF(CCAGAACGGATTCACGGC), ptxAR (CTGCTGCTGGTGGAGACGA), bvgAF (AGGTCATCAATGCCGCCA), bvgAR (GCAGGACGGTCAGTTCGC), fhaBF (CAAGGGCGGCAAGGTGA), fhaBR (ACAGGATGGCGAACAGGCT), rpoBF (GCTGGGACCCGAGGAAAT), rpoBR (CGCCAATGTAGACGATGCC). BP2497 were designed in a previous study: BP2497F (TCGGATCGCACCAATTACTTC) and BP2497R (CCTTGGCGATCAGCGAGTT) (10). The following primers were used for *Gapdh:* gapdhf (CATGGCCTTCCGTGTTCCT) and gapdhr (GCGGCACGTCAGATCCA).

## Results and Discussion

### Establishment of a murine model to perform *in vivo* transcriptomic analysis of *B. pertussis*

The main objective of this study was to perform RNA sequencing on *B. pertussis* during lung infection. Our previous studies with the respiratory pathogen *P. aeruginosa* demonstrated the feasibility of sequencing both bacterial and host pathogen during infection. However, they also highlighted the need to use high bacterial loads (10^8^ to 10^9^ CFU/organ) to obtain sufficient RNA for purification and analysis. In addition, our previous attempts to perform RNAseq from CD1 mouse trachea yielded very few reads of bacterial RNA, demonstrating the challenge of these types of workflows. Therefore, in this work, we first optimized the murine model of pertussis to maximize the recovery of bacteria during infection. To this end, we performed intranasal infections of outbred CD1 mice (traditional model used for vaccine development) and in immunodeficient NSG mice. We hypothesized that immunodeficient mice would be unable to clear bacterial infection and have higher bacterial burdens in the airways which would increase the recovery rate of bacterial RNA for sequencing. Mice were infected with 10^7^ CFU/mouse (CD1) or 4x10^6^ CFU/mouse (NSG) and euthanized 24 or 72 hr post infection. Bacterial burden was determined in the nasal passages by performing a nasal wash, and in the trachea and lung by generating tissue homogenates. As expected, significantly higher bacterial burden was observed in the nasal wash, trachea, and lung of NSG mice compared to CD1 mice (Fig. 1). The bacterial burden in the nasal wash and lung remained constant between day 1 and 3 in CD1 mice, and even significantly decreased in the trachea (Fig. 1). However, bacterial burden significantly increased over time in NSG infected mice in both the upper and the lower respiratory airways (Fig 1). Overall, the lung samples of NSG mice at the later time point had the highest bacterial burden. Therefore, for subsequent experiment, the lungs from NSG mice infected with *B. pertussis* strain UT25 were harvested two days post infection and used for RNA preparation for NGS.

**Figure 1.**
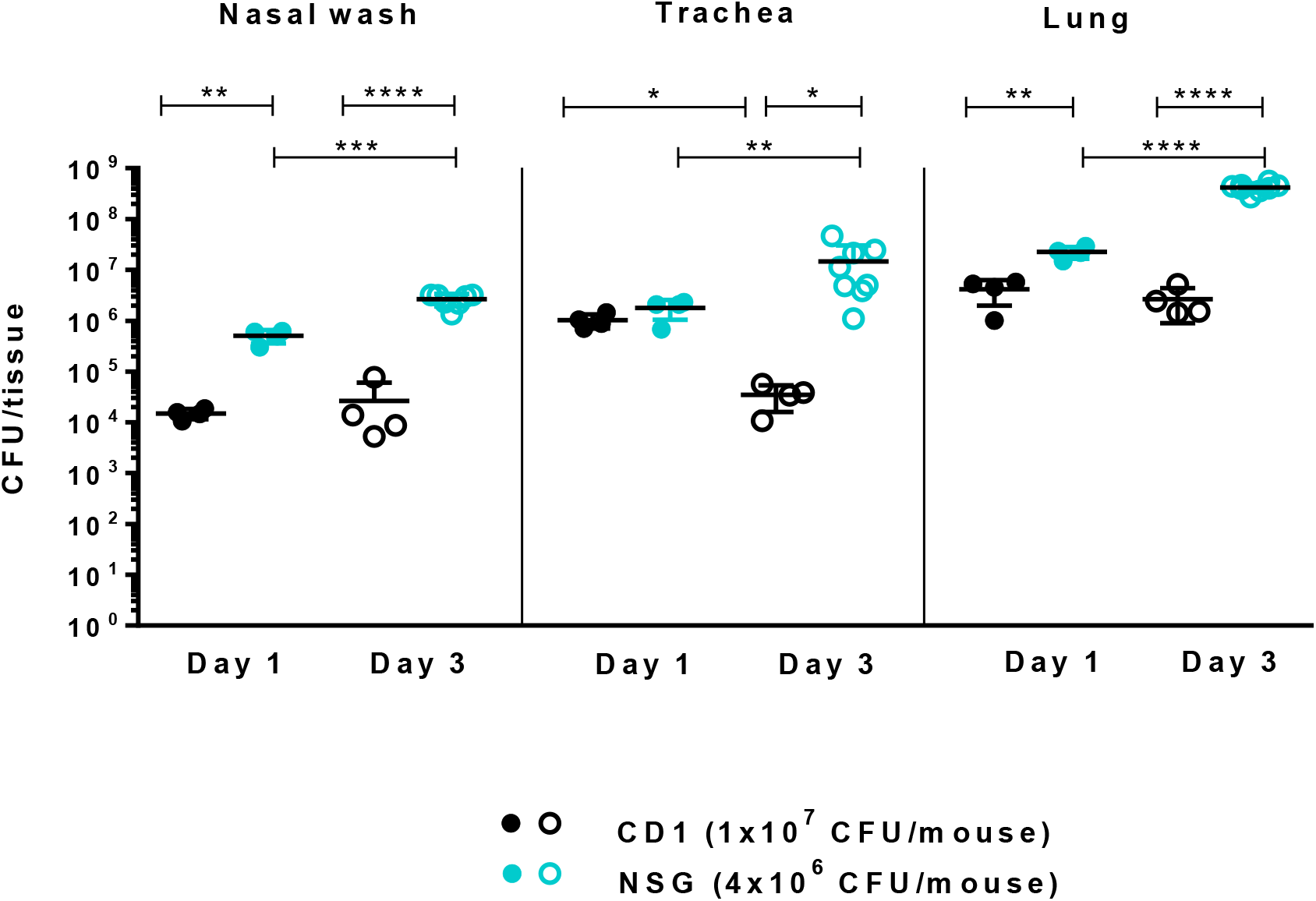
*B. pertussis* burden in outbred CD1 mice compared to highly sensitive NSG mice. NSG mice were infected with 4x10^6^ CFU and CD1 mice were infected with 2x10^7^ CFU of the *B. pertussis* strains UT25. At 1 and 3 days post infection the bacterial burdens in the nasal wash, trachea, and lungs were determined. Four to eight mice were used for each time-point. Groups were compared independently using a t-test with Tukey correction (* *p*<0.05; ** *p*<0.01; *** *p*<0.001; **** *p*<0.0001).

#### RNAseq analysis of NSG lungs infected with *B. pertussis*

To perform dual RNAseq (analyze both host and pathogen), RNA was isolated from the infected NSG lungs per the standard RNeasy column based kit. Illumina Scriptseq libraries were prepared and sequenced on a Hiseq 1500. 100 million 2x50bp reads were devoted to each sample which contained 3 pooled *B. pertussis* infected NSG lungs. Approximately 200,000 reads mapped to the *B. pertussis* Tohama I reference genome for each sample. Upon inspection of the reads, it was apparent that the mouse RNA of the sample was cross mapping to the bacterial genome. This non-specific cross-mapping was due to sequence similarities between the bacterial and the murine genomes. In order to focus on only true *B. pertussis* reads, we extracted only the reads that mapped with 100% identity in pairs. This removed the false positive mappings as observed in Fig. 2. However, this procedure also significantly decreased the number of *B. pertussis* reads. Therefore, we sampled the *in vitro* UT25 reads down to 15,000 to allow for a more accurate comparison to the NSG mouse *B. pertussis* reads. Overall we observed that only 0.014% of the total reads corresponded to the transcriptome of *B. pertussis*, which is insufficient for high quality transcriptome analysis. The murine reads greatly outnumbered the pathogen reads (Table 1). We were able to detect up to 828 genes with at least 1 true *B. pertussis* read. When we compared the *in vitro* and *in vivo B. pertussis* reads (Table 2), we observed promising data suggesting that the changes in gene expression observed in these samples are relevant for the pathogenesis of *B. pertussis*. We performed String analysis to better understand the associations between the genes differentially regulated between the two datasets (Fig. 3). In this analysis, we observed the alcaligin siderophore biosynthesis genes in the infected NSG lungs (Table 2 and Fig. 3A). Alcaligin is a siderophore that sequesters iron for the pathogen and is an important factor required for infection. We also detected *fhaB* in the infected lung tissue. FhaB is an adhesin that allows *B. pertussis* to adhere to the airway during infection. In previous experiments we observed that *fhaB* is one of the most abundant transcripts in the transcriptome (10). We also detected both pertussis and adenylate cyclase toxin transcripts in the infected lung tissue (Table 2). Overall in this limited analysis, *B. pertussis* gene expression was higher *in vitro*. It is likely that *B. pertussis* grows faster *in vitro* and is transcriptionally more active (Fig. 3B). From this analysis, we concluded that method optimization to enrich the RNA of *B. pertussis* was required in order to characterize the *in vivo* transcriptome of the bacterium in the mouse.

**Figure 2.**
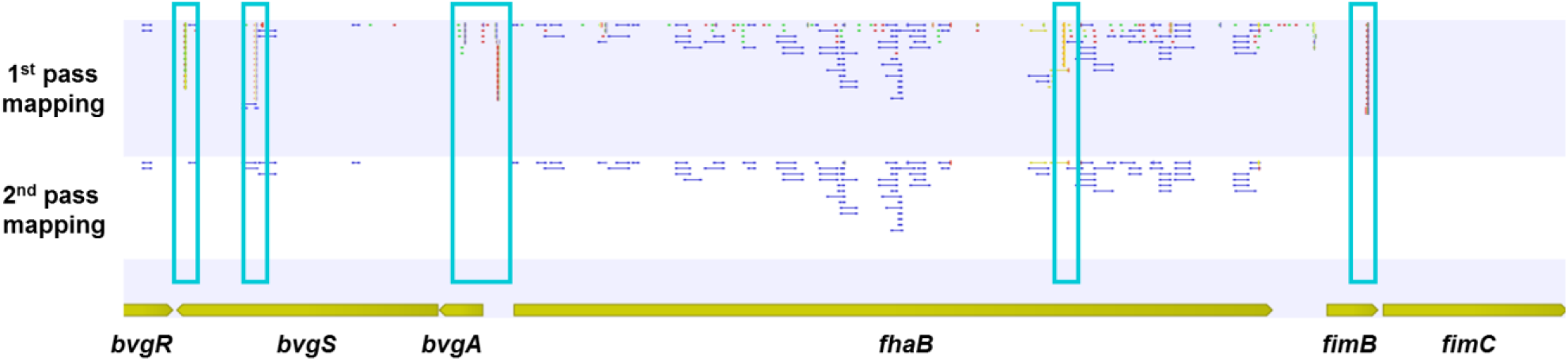
Visualization of the *bvgRSA* and *fhaB* loci mapped reads. Reads were mapped to the *B. pertussis* genome (1^st^ pass mapping) and cross-mapping murine RNA was observed. The stringency of identify was increased (2^nd^ pass mapping) and only paired reads were considered. In the blue boxes, examples of cross mapping reads are shown.

**Figure 3.**
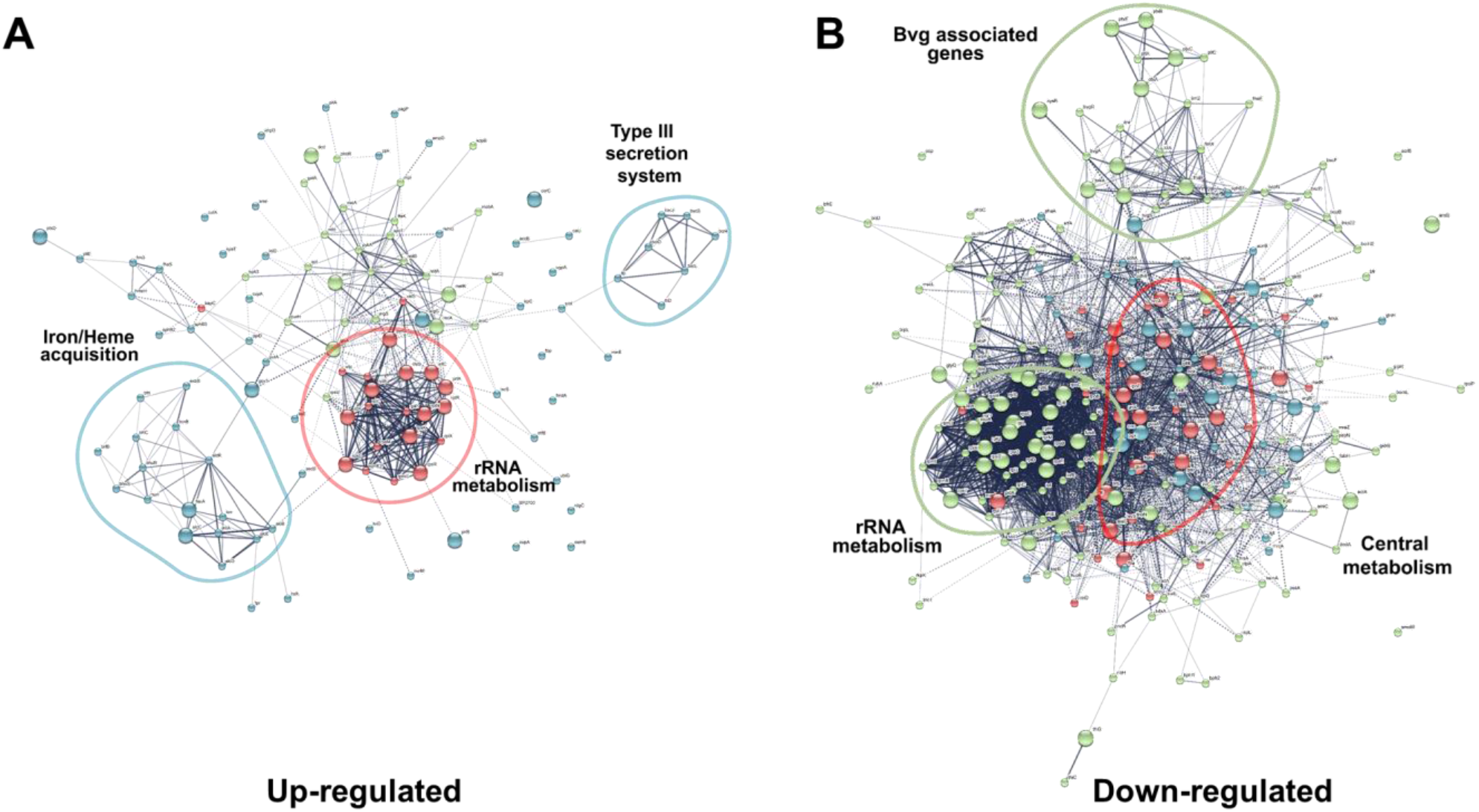
String analysis of differentially regulated genes during infection. Up- (A) and down-regulated (B) annotated *B. pertussis* genes in the lung compared to *in vitro* were analyzed and represented using the String database (REF) and kmeans clustering.

**Table 1.** Summary of RNA-seq gene reads and mapping.

| Sample | Total reads in sample | % of reads mapped to *B. pertussis* genome | Reads mapped to *B. pertussis* genome | Number of genes with 10 reads | Number of genes with 1 read | % of the genes in the genome with at least 1 read |
| --- | --- | --- | --- | --- | --- | --- |
| NSG mouse lung infected with UT25 | 109,878,438 | 0.01365 | 14,995 | 69 | 828 | 21.47 |
| *B. pertussis* UT25 growing in SSM | 15,000 | 97.35 | 14,602 | 94 | 1,343 | 34.48 |

**Table 2.**

| Gene | gene product | Fold change | SSM reads | SSM RPKM | NSG reads | NSG RPKM |
| --- | --- | --- | --- | --- | --- | --- |
| **Genes of interest and virulence factors** |  |  |  |  |  |  |
| *alcA* | Alcaligin biosynthesis enzyme | ∞ | 0 | 0 | 27 | 1256 |
| *alcB* | Alcaligin biosynthesis protein | ∞ | 0 | 0 | 16 | 1702 |
| *alcC* | Alcaligin biosynthesis protein | ∞ | 0 | 0 | 12 | 417 |
| *alcE* | Putative iron-sulfur protein | ∞ | 0 | 0 | 1 | 80 |
| *bcr4* | Siderophore [Alcaligin] translocase AlcS | 11 | 2 | 525 | 47 | 5804 |
| *bcrD* | Type III secretion inner membrane channel Protein (LcrD,HrcV,EscV,SsaV) | -2 | 8 | 491 | 9 | 260 |
| *bfrC* | Ferrichrome-iron receptor | ∞ | 0 | 0 | 12 | 353 |
| *bfrI* | Ferrichrome-iron receptor | ∞ | 0 | 0 | 12 | 367 |
| *brkA* | Serum resistance protein | -10 | 23 | 1039 | 5 | 106 |
| *bvgA* | Virulence factors transcription regulator | -17 | 8 | 1739 | 1 | 102 |
| *bvgS* | Hybrid sensory histidine kinase | -20 | 19 | 700 | 2 | 35 |
| *cyaA* | Hemolysin-adenylate cyclase precursor | -3 | 13 | 348 | 11 | 138 |
| *fhaB* | Filamentous hemagglutinin/adhesin | -7 | 275 | 3496 | 83 | 497 |
| *fhaL* | Adhesin | -4 | 9 | 98 | 5 | 26 |
| *fhaS* | Adhesin | 6 | 1 | 18 | 12 | 101 |
| *fim2* | Serotype 2 fimbrial subunit precursor | 1 | 1 | 216 | 3 | 306 |
| *fim3* | Serotype 3 fimbrial subunit precursor | 3 | 17 | 3786 | 96 | 10063 |
| *groEL* | Heat shock protein 60 family chaperone GroEL | -3 | 67 | 5582 | 45 | 1765 |
| *ompA* | Outer membrane protein A precursor | -28 | 66 | 15532 | 5 | 554 |
| *prn* | Pertactin precursor |  |  |  |  |  |
| *ptxA* | Pertussis toxin subunit 1 precursor | -3 | 16 | 2706 | 11 | 875 |
| *ptxB* | Pertussis toxin subunit 2 precursor | -6 | 12 | 2414 | 4 | 379 |
| *ptxC* | Pertussis toxin subunit 3 precursor | -2 | 10 | 2002 | 14 | 1319 |
| *ptxD* | Pertussis toxin subunit 4 precursor | 1 | 5 | 1492 | 13 | 1826 |
| *ptxE* | Pertussis toxin subunit 5 precursor | -11 | 16 | 802 | 3 | 71 |
| *ssrA* | Transfer-messenger RNA (tmRNA) | -20 | 1094 | 388193 | 114 | 19039 |
| *tcfA* | Tracheal colonization factor precursor | -7 | 38 | 2677 | 11 | 365 |
| *tuf_1* | Translation elongation factor Tu | -1 | 22 | 2530 | 33 | 1786 |
| *vag8* | Autotransporter | -5 | 46 | 2293 | 19 | 446 |
| **rRNAs** |  |  |  |  |  |  |
| *BPr01* | SSU rRNA | 82 | 17 | 1520 | 2959 | 124514 |
| *BPr02* | LSU rRNA | 35 | 81 | 3850 | 6103 | 136515 |
| *BPr05* | LSU rRNA | 4 | 77 | 3659 | 638 | 14271 |
| *BPr06* | SSU rRNA | 15 | 18 | 1609 | 573 | 24112 |
| *BPr08* | LSU rRNA | 3 | 80 | 3802 | 583 | 13041 |
| *BPr09* | SSU rRNA | 17 | 17 | 1520 | 596 | 25080 |
| *BPs02* |  | -14 | 285 | 94289 | 42 | 6540 |
| **Hypothetical genes** |  |  |  |  |  |  |
| *BP0168* | Putative membrane protein | 14 | 1 | 152 | 30 | 2142 |
| *BP0840* | Putative membrane porin protein precursor | -46 | 303 | 35839 | 14 | 779 |
| *BP2374* | Hypothetical gene | ∞ | 0 | 0 | 42 | 13675 |
| *BP2410* | Hypothetical gene | 19 | 1 | 102 | 40 | 1926 |
| *BP3084* | Putative metal chaperone, Zn homeostasis | 3 | 10 | 1227 | 61 | 3524 |
| *BP3402* | Zinc metalloprotease YfgC precursor | ∞ | 0 | 0 | 94 | 3992 |

#### Method optimization for the extraction of bacterial RNA using differential filtration

Despite murine model optimization to maximize bacterial RNA recovery the total number of reads from bacterial RNA remained low. To increase bacterial sequence coverage without rising sequencing cost by increasing the total number of sequencing lanes and total reads, we developed a novel methodology to separate bacterial from eukaryotic RNA. This method is based on the difference in size between bacterial and eukaryotic cells present in lung homogenates. We first performed method optimization using *in vitro* grown bacterial samples of *E. coli, P. aeruginosa, B. bronchiseptica*, and *B. pertussis. In vitro* grown cultures of various bacterial pathogens were filtered through a 5 μm filter, and plated before and after filtration to estimate percentage recovery of the bacteria. Overall, 30 to 60% of the bacteria present in the suspension were recovered after filtration (Figure 4). Recovery rate was different between the species, with *B. bronchiseptica* showing the highest recovery rate and *P. aeruginosa* the lowest. These differences are potentially due to the variation in size of these bacteria, *P. aeruginosa* cells measuring up to 3μm in length, while *Bordetellae* cells measure less than 0.7 μm in length. From these results, we concluded that 5 μm filters could be used to recover a significant number of bacteria from suspensions.

**Figure 4.**
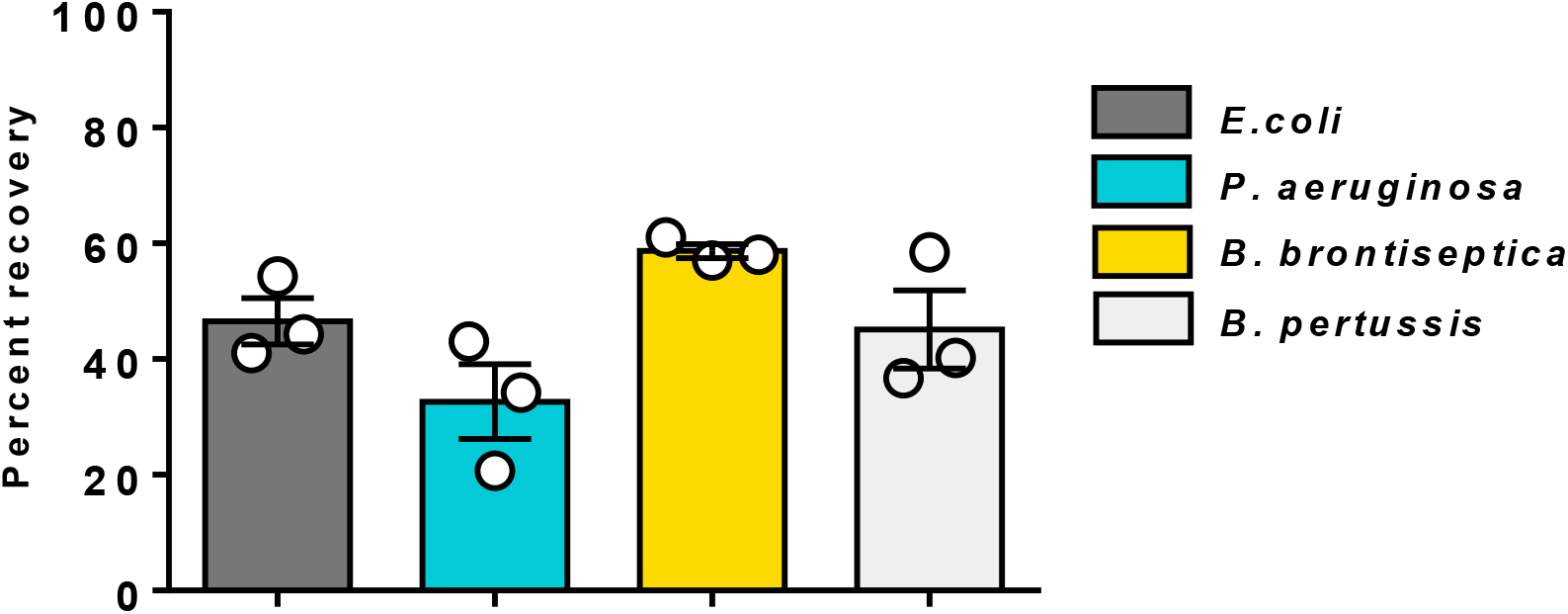
Recovery of bacteria after syringe filtration through a 5 μm filter. Percent recovery of the bacteria present in a suspension prepared from *in vitro* grown cultures after filtration with a 5 μm filter. Each bar shows the mean percentage recovery of bacteria from suspensions at densities ranging from OD_600_ 0.3 to OD_600_ 0.003 to illustrate that the recovery rate does not depend on bacterial suspension density but on bacterial size.

#### Method optimization for the extraction of bacterial RNA from infected tissue

In the next phase of our protocol development, we applied filtration to homogenized tissue to separate bacterial cells from the rest of the homogenate. Infected tissues were first gently homogenized using a mechanical Dounce glass homogenizer to separate the cells in the lung while causing minimal cell lysis compared to enzymatic digestion. Homogenates were then filtered through a 70 μm filter to remove large clumps and debris. The resulting filtrate was pushed through a 5 μm filter to remove eukaryotic cells. This second filtrate, containing bacteria and residual eukaryotic DNA released from cell lysis during homogenization, was then centrifuged to pellet bacteria. The bacterial pellet was finally re-suspended in RNA protect and processed for extraction (Fig. 5). Bacterial loads in nasal wash, trachea and lung were quantified in the tissue homogenate and in the final bacterial pellet after filtration (Fig. 6). From these data, we estimated the range of viable bacteria needed for RNAseq analysis. We estimate that RNA from 10^4^ *B. pertussis* CFU is the minimum that can be used for *in vivo* seq (assuming 100% of the sample is pathogen). This estimation is based on our filtration method and RNA purification using RNAsnap, and assumes that the RNA obtained is DNA-free. For samples in which these yields cannot be achieved, low input kits such as the Nugen Ovation RNAseq V2 library preparation kit can be used to perform the analysis with picogram amounts of RNA.

**Figure 5.**
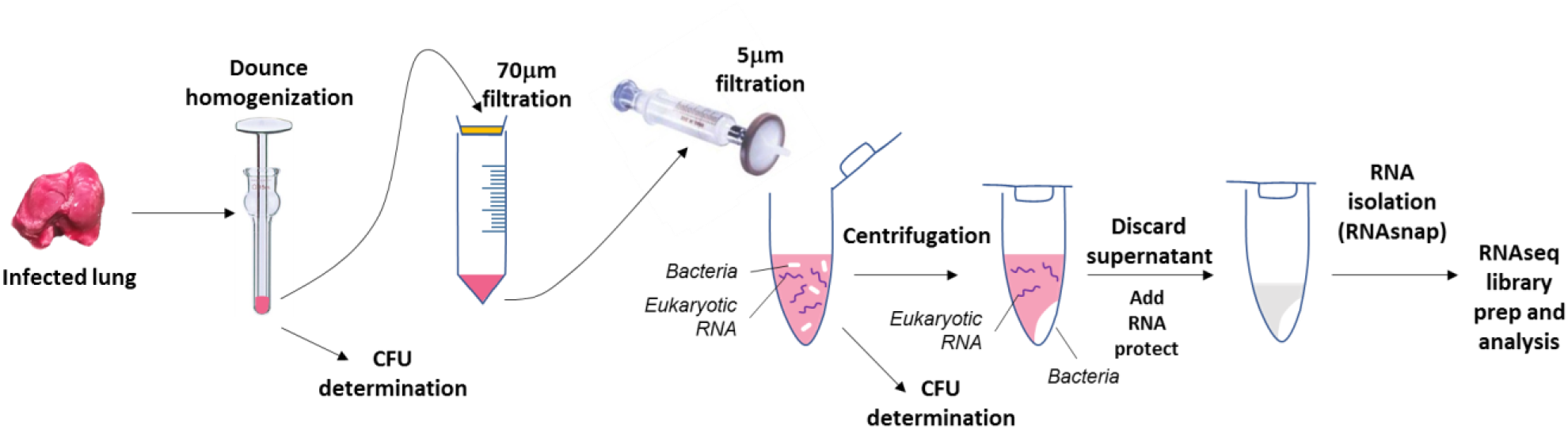
Isolation of *B. pertussis* from infected murine lung for preparation of RNA for *in vivo* RNAseq. Samples are homogenized using a Dounce glass homogenizer and tissue debris are removed by filtration through a 70 μm filter. Eukaryotic cells are excluded by 5 μm filtration. Bacteria are isolated by centrifugation and contaminating murine RNA (due to lysis of cells) is removed. Bacterial cells are stabilized by RNAprotect until RNAsnap RNA isolation.

**Figure 6.**
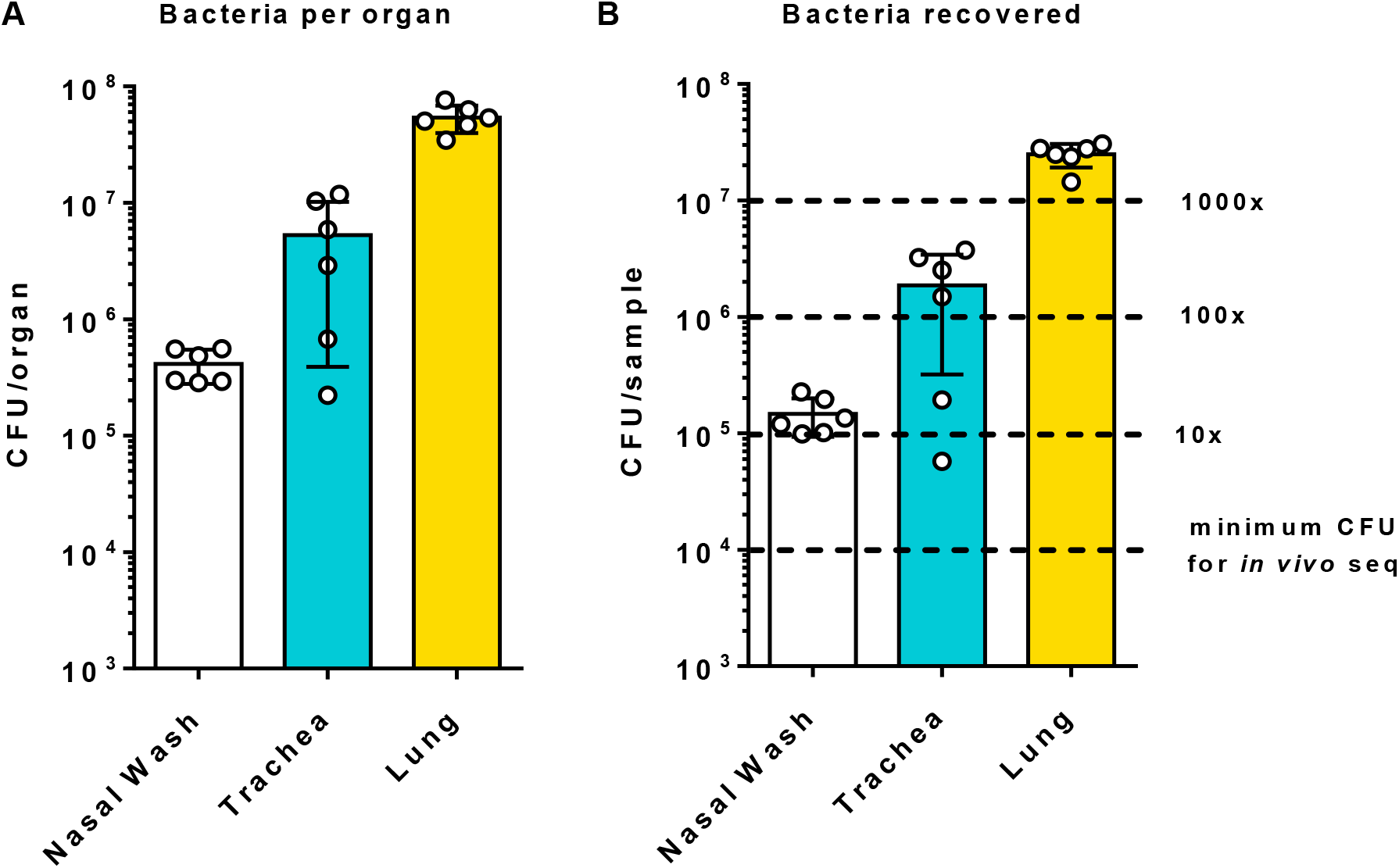
Isolation of *B. pertussis* from infected NSG mouse lungs. NSG mice were infected with *B. pertussis* strain UT25. At two days post infection, the tissues were collected and the number of input and output (post filtration) were quantified. A. Bacterial load in each tissue after homogenization. B. Number of bacteria recovered from each tissue after implementing the filtration strategy described in Fig. 5. The estimated number of bacteria needed to perform RNAseq assuming 100% bacteria RNA in the sample are shown using dashed lines.

#### qPCR estimation of *rpoB* to *Gapdh* transcript levels

To sequence the transcriptome of *B. pertussis* during infection, it is important to have a high bacteria to host RNA ratio in the sample. To estimate this ratio, we performed absolute qPCR to count the numbers of transcripts in cDNA samples from the *B. pertussis* infected NSG mice (Fig. 7). We added this quality control metric to select samples with high amounts of bacterial RNA that have the best probability of successful RNAseq analysis. Standard curves of *rpoB* and *Gapdh* templates were generated and used to calculate *rpoB* and *Gapdh* transcript copy numbers. We observed that not all samples had the same pathogen to host copy ratios (Fig. 7). However, the data suggested that in most samples there was a 1:4 ratio of *rpoB* to *Gapdh*. We also filtered a *B. pertussis* infected CD1 lung and observed a similar *rpoB* to *Gapdh* ratio. This promising data suggests that we can obtain sufficient RNA, even from mice that are not highly susceptible.

**Figure 7.**
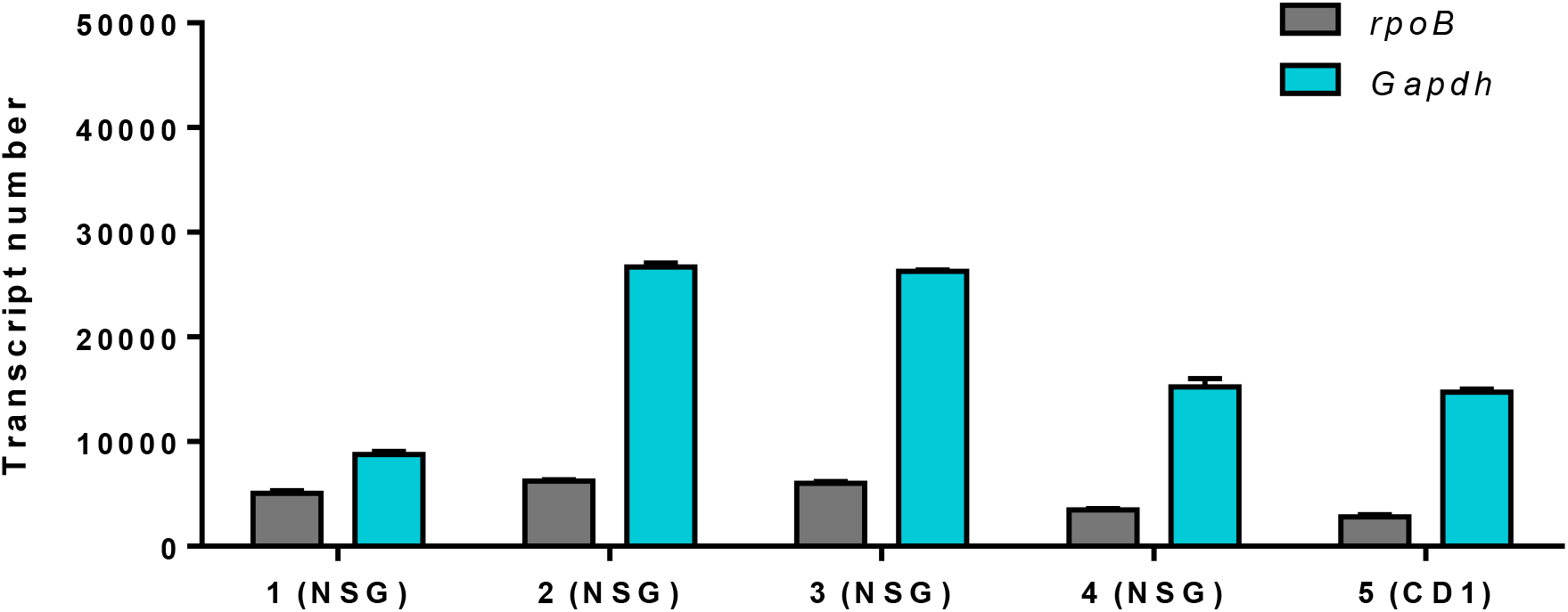
Absolute quantification of *rpoB* and *Gapdh* transcripts in the filtered *B. pertussis* infected NSG lungs. Standard curves of known amounts of *rpoB* or *Gapdh* targets were established and used to calculate the number of transcripts of each gene in cDNA from the lung of *B. pertussis* infected NSG mice. Four NSG mice were analyzed along with one outbred CD1 mouse. The CFU present in the lung of the NSG mice are shown in Figure 6A. The CD1 mouse had 10^7^ viable *B. pertussis* when the lung was harvested. These absolute qPCR methods also provides a quality control metric to validate samples before proceeding to RNAseq library preparation.

#### qRT-PCR analysis of *in vitro* and *in vivo* grown *B. pertussis*

To confirm that this method could allow detection of bacterial transcript during infection, we performed qRT-PCR analysis on *B. pertussis* RNA samples proceeding from SSM *in vitro* cultures and RNA from *B. pertussis* infected NSG lung samples (Fig. 6 and 8). We selected the *cya* and *ptx* genes that encode the major toxins as well as *fhaB, bvgA*, and *BP2497*. We hypothesized that the expression of the genes encoding these toxins would be more highly expressed in the mouse compared to *in vitro*. We observed increased *cya* and *ptx* gene expression in all of our NSG mice (Fig. 8). Expression of *fhaB*, the major adhesin, was decreased *in vivo* compared to *in vitro*. The genes *cya* and *ptx* are classified as “class II Bvg system genes” (3) whereas *fhaB* is a “class I gene”. Class I genes are usually expressed at different levels than class II genes, which could explain why *cya, ptx* and *fhaB* are differentially regulated. In addition, it is possible that the bacteria tightly adhered to epithelial cells during infection are not collected using this purification method, which would introduce bias and explain differences in gene expression involved in bacterial adhesion. This caveat will be further investigated as the project moves forward. At this phase, we are most interested in successfully performing the feasibility of performing *in vivo* RNAseq on *B. pertussis* in the murine lung.

**Figure 8.**
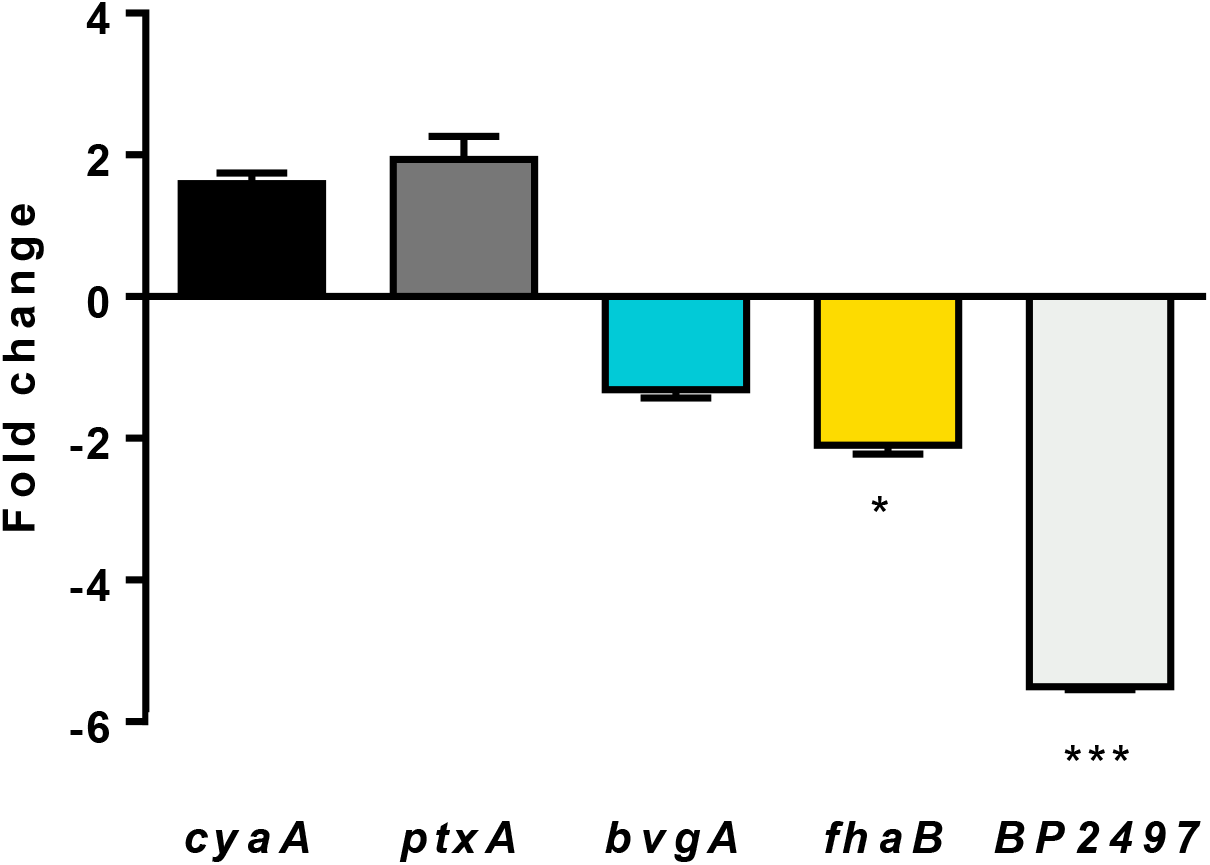
qRT-PCR analysis of *B. pertussis* growing *in vitro* compared to *in vivo* in the NSG lung. *B. pertussis* strain UT25 was grown in SSM liquid media for control comparison. The NSG lung samples prepared by the filtration protocol were compared to the SSM control data. Genes encoding the adenylate cyclase toxin (*cya*) and the pertussis toxin (*ptxA*) were increased *in vivo* compared to *in vitro*. The *bvgA* and *fhaA* genes were slightly decreased in expression. *BP2497* encodes a putative protease and it was also decreased in the murine lung. Data from three biological replicates with three technical replicates each. Standard deviations are calculated from variations between biological replicates and compared using a *t*-test (* *p*<0.05; *** *p*<0.001).

## Conclusions

In this report, we have detailed our progress on developing RNAseq workflows to characterize the transcriptome of *B. pertussis* infecting the murine lung. Bacterial pathogens are small yet can cause substantial harm to the host. *B. pertussis* releases toxins which block or suppress both the innate and adaptive immune responses. These toxins are essential to infection but little is known about the roles of the majority of the genes encoded in the genome of *B. pertussis*. We hypothesize that the *B. pertussis* transcriptome *in vivo* will be significantly different than the *in vitro* transcriptome in media such as on BG or SSM. It is possible genes that are highly expressed during infection could be exploited as vaccine antigens. By answering the simple question of “what does the pathogen express when it infects?”, we believe that significant advancement can be made which will potentially impact vaccines and treatments. Our initial attempts showed that performing dual-seq on a pathogen that is both growing slowly and is present in at low levels in tissue samples is challenging. These technical difficulties led us to re-formulate our ideas and come up with an innovate strategy. By combining a novel filtration method to separate the bacteria from host tissue and exhaustive qRT-PCR quality controls, we have obtained RNA with a pathogen to host ration high enough to perform RNAseq in the near future. Next generation sequencing applications such as RNAseq have greatly expanded over the past few years but we believe there is still a need to further develop RNA preparation workflows to truly take advantage of these technologies. There are many other bacterial pathogens for this type of methodology could potentially be applied. We envision that this strategy could also be used in other models systems such as the baboon model of pertussis. We hypothesize that depending on the lower limits of detection, it may also be possible to isolate pathogens directly from human patients and determine the “human specific” transcriptomes of the pathogen. Our current goal is to fully characterize the transcriptome of the human pathogen *B. pertussis* and we will also seek to apply this strategy to other pathogens and extend our knowledge of how pathogens infect.

## Acknowledgements

*B. pertussis* strains UT25 was kindly provided by Dr. Sandra Armstrong (University of Minnesota). We would like to thank the WVU Transgenic Animal Core Facility for providing the NSG mice for this study.

## Author contributions

T.W. and J.H. performed murine experiments, prepared RNA, validated RNA for RNAseq, performed qRT-PCR, analyzed data, and composed the manuscript. D.T.B performed murine experiments analyzed data, and composed the manuscript. M.B. designed / performed murine experiments, analyzed data, and composed the manuscript. F.H.D designed / performed experiments (murine infection model, RNAseq, and etc), analyzed data, and composed the manuscript.

